# Impacts of dams on freshwater turtles: a global review to identify conservation solutions

**DOI:** 10.1101/2021.10.21.465338

**Authors:** Andrea Bárcenas-García, Fernanda Michalski, William H. Morgan, Rebecca K. Smith, William J. Sutherland, James P. Gibbs, Darren Norris

## Abstract

**Background and Research Aims:** Dams impact freshwater ecosystems and biodiversity. Freshwater turtles are at direct and indirect risk due to changes caused by damming including the loss of terrestrial and aquatic nesting habitats, changes to food availability and blocking movement. Effective management of these impacts requires robust evidence in order to gain an understanding of conservation solutions that work.

**Methods:** We reviewed the global scientific literature that evaluated the impact of dams on freshwater turtles, and carried out additional searches of literature published in seventeen languages for studies evaluating actions to mitigate dam impacts.

**Results:** The search produced 47 published articles documenting dam impacts on 30 freshwater turtle species from seven families (Chelidae, Chelydridae, Emydidae, Geoemydidae, Kinosternidae, Podocnemididae and Trionychidae) in 13 countries. Few studies were found from Europe and Asia and none from Africa. Most studies were from temperate latitudes, where studies focused more on adults and less threatened species compared with tropical latitudes. More than half of the studies (57%, n = 27) suggested actions to help mitigate dam impacts. Yet, only five studies (three temperate and two tropical) documented the effect of interventions (dam removal, flow management, artificial pond maintenance and community-based action).

**Conclusion:** These findings demonstrate a serious lack of documented evidence evaluating mitigation actions for dam impacts on freshwater turtles. Implications for Conservation: This lack of evidence reinforces the importance of strengthening and maintaining robust long-term studies needed to develop effective and adaptive conservation actions for this group of threatened vertebrates particularly in tropical regions.

## Introduction

Biodiversity declines are more accelerated in freshwater compared to marine or terrestrial ecosystems (Harrison et al., 2018; He et al., 2018). Freshwater environments provide a variety of natural resources and have been subject to intense human management for millennia (Fitzhugh & Richter, 2004; Pradinaud et al., 2019). Structures including dams and locks are used to manage flows and provide storage to meet myriad needs of expanding human populations (e.g. drinking water, agricultural irrigation, hydropower and transport) and today nearly half of all rivers have been modified by dam construction (Grill et al., 2019). Dams are considered primary threats to freshwater species, as well as the surrounding ecosystems including floodplains, wetlands and riparian habitats (Harper et al., 2021; Zarfl et al., 2019). Although populations of freshwater vertebrates have declined at more than twice the rate of terrestrial or marine vertebrates (Grooten & Almond, 2018; Tickner et al., 2020), relatively few studies have evaluated the impact of dams on vertebrates (dos Santos et al., 2021; He et al., 2018).

Dams and associated up- and downstream fragmentation and flow regulation contribute to the loss of river connectivity and freshwater biodiversity (Grill et al., 2019; Harper et al., 2021). Species that inhabit freshwater ecosystems are vulnerable to extinction due to dams impacts (Tickner et al., 2020), as their life histories and critical habitats often strongly depend on the hydrological regime (Zarfl et al., 2019). Most studies have focused on dam impacts to fish populations because fishes are often both an important source of protein as well as having high commercial and recreational value. The relative lack of studies on other vertebrate fauna is surprising considering that damming could contribute to the extinction e.g. dolphins (Brownell Robert et al., 2017; Turvey et al., 2010) or extirpation of diverse vertebrate species in impacted basins e.g. turtles (Jian et al., 2013; Santoro et al., 2020). Despite the known impacts, there is little available evidence documenting dam mitigation interventions for aquatic fauna such as freshwater turtles (CEE, 2021; dos Santos et al., 2021; Sainsbury et al., 2021; Tickner et al., 2020).

Turtles are an ancient, widespread and instantly recognizable group that not only provide highly valued cultural, medicinal and economic resources across the globe (Haitao et al., 2008; Liu et al., 2020; Lovich et al., 2018; Mendiratta et al., 2017; Sigouin et al., 2017; TTWG et al., 2017) but also provide inspiration for the development of 21^st^ century biomimetic robotics (Kim et al., 2012; Soliman et al., 2021). Although turtles are important to both humans (Stanford et al., 2020) and aquatic ecosystem functioning (Lovich et al., 2018; Moll & Moll, 2004b) they are the most threatened group of freshwater vertebrates (Rhodin et al., 2018; Stanford et al., 2020; Tickner et al., 2020). Even protected areas are insufficient to buffer freshwater turtles from human impacts (Howell et al., 2019; Norris et al., 2019). The meat and eggs of freshwater turtles are widely used as food resources (Stanford et al., 2020; TCC, 2018), while the fat, viscera and shell are also used e.g. in traditional medicine (Dudgeon, 2019; Pezzuti et al., 2010). Dams have also been identified as a major threat for many freshwater turtle species (Bodie, 2001; Moll & Moll, 2004a), including for 11 of the 25 most threatened tortoise and freshwater turtle species (TCC, 2018). For example, planned hydropower dams may permanently flood 73% of potential nesting habitat of the Yangtze giant softshell turtle (*Rafetus swinhoei)* the rarest freshwater turtle in the world (Jian et al., 2013; TCC, 2018). This loss of habitat, coupled with the historic exploitation of *R. swinhoei* throughout its range, contributes to increasing extinction risk (Jian et al., 2013; Stanford et al., 2020). Previous studies recognize dams as an indirect threat to freshwater turtles (Moll & Moll, 2004a), however, more recent reports provide evidence that as barriers to movement dams directly cause mortality in adult turtles e.g. males and females of the aquatic yellow-spotted river turtles (*Podocnemis unifilis)* that are obliged to move overland around the Belo Monte dam complex in Brazil and become trapped, overheated and/or dehydrated (JGP Consultoria, 2019). Despite widespread impacts, studies of freshwater turtle population dynamics remain scarce, as there is a lack of robust information on the life history of many species particularly those found in the tropics (Rachmansah et al., 2020; Rhodin et al., 2018; TCC, 2018). As such, the ecological requirements and life history of at least 30% of turtle species are as yet unknown, making their conservation status difficult to evaluate (Rhodin et al., 2018).

The continued expansion of damming across the globe requires an evaluation of the available evidence for the development of effective mitigation actions. In this paper we synthesize studies that have evaluated the conservation of freshwater turtles in areas around the world altered by dams. Our aim was to identify research trends, gaps in current knowledge, mitigation actions both proposed and tested about dam impacts, and conservation solutions for freshwater turtles.

## Material and methods

### Literature search

Complementary approaches were adopted to identify not only threats of dams on freshwater turtles but also the conservation solutions to minimize and mitigate impacts. A review of the scientific literature following the protocol of Preferred Reporting Items for Systematic Reviews and Meta-Analyses [PRISMA (Page et al., 2021)] was conducted in the ISI Web of Science (Core Collection) database (Fig. 1). Web of Science searches were conducted on 13 October 2021 and updated on 6 April 2022 to include articles published from 1945 to 1 April 2022. A search of all Web of Science database fields included the following combination of English terms: (turtle* OR terrapin* OR Chelon* OR Testudines OR Cryptodira OR Pleurodira) AND (hydropower OR dam* OR hydroelectric* OR reservoir*). Conference proceedings were not included in the search.

**Figure 1.**
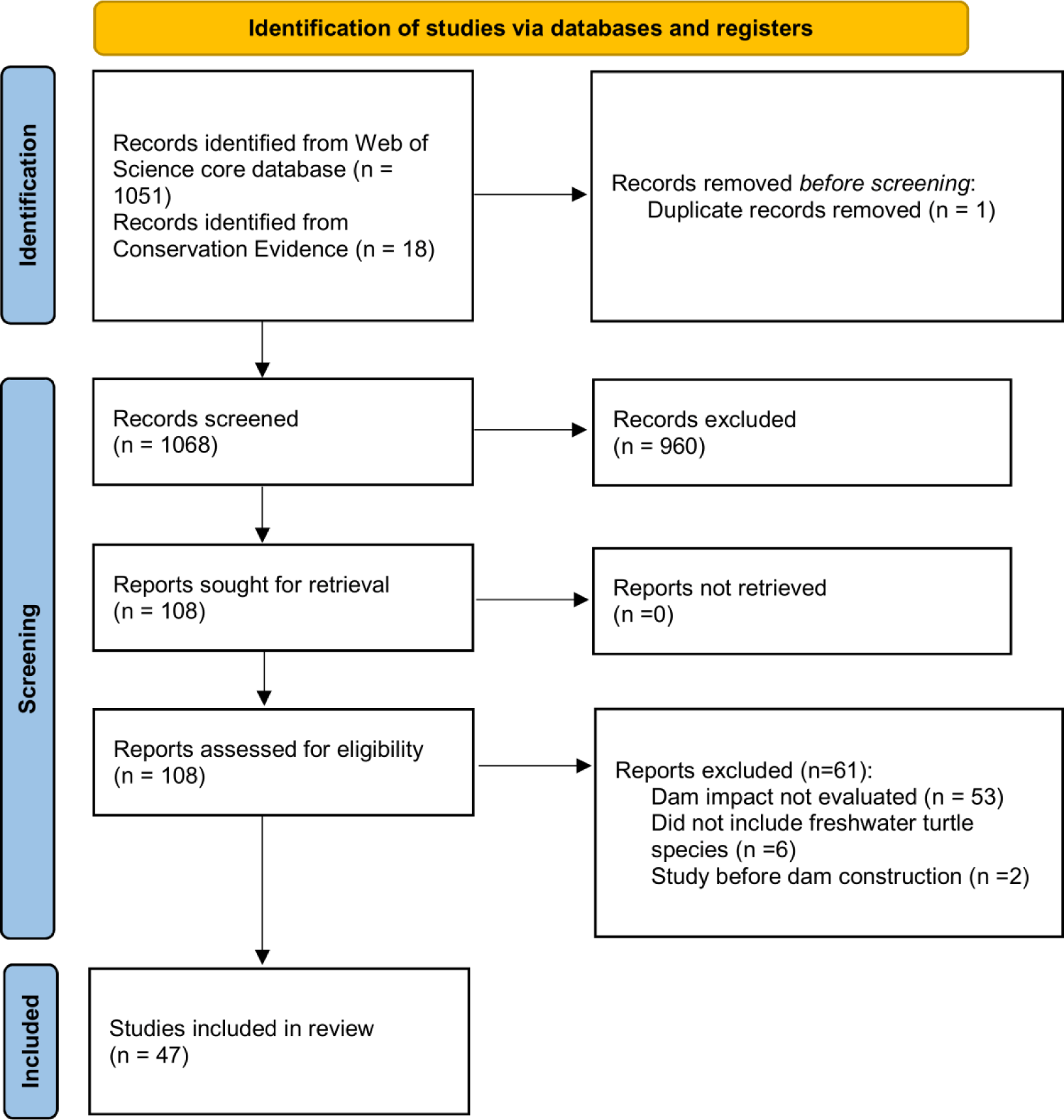
Literature search. Flow diagram with the Preferred Reporting Items for Systematic Reviews and Meta-Analyses (PRISMA) process steps and number of studies excluded and included.

### Selection criteria and process

The Web of Science searches identified a total of 1051 articles (Fig. 1, Supplemental material S1, data available at doi: 10.17605/OSF.IO/KQ573). All article titles and abstracts were read and screened independently by two authors (AB, DN) to retain studies that potentially included freshwater turtles and dams (Gough et al., 2020). Screening results were combined retaining all those identified by either author as potentially including freshwater turtles and dams. A subsample of 106 articles (10% of the total) was also screened by a third author (WM) to evaluate the combined screening result (Collaboration for Environmental Evidence, 2013). As the screening conducted by the third author was in 100% agreement with the combined result, no additional adjustments to the screening process were taken. The full text of 108 articles that passed screening was then read and articles were assessed based on two criteria: 1) the study had to include data on at least one freshwater turtle species; 2) the study measured current- or post-construction effects/impacts and/or mitigation actions of dams (including removal). We included original research articles with primary and secondary data, including field based, modelling inference, interviews and laboratory (e.g. genetic) studies. Studies that included only summarized versions of compiled primary data (e.g. reviews and perspectives) were excluded. Studies that evaluated river channel alternations not associated with dams (e.g. channel widening (Usuda et al., 2012)) or where dam impacts were discussed based on unconfirmed secondary narratives lacking methodological details (Kitimasak et al., 2005) and/or merely discussed (e.g. (Tornabene et al., 2017)) were also not included. This approach was adopted to enable us to establish the most robust evidence possible of directionality for all reported impacts.

### Conservation Evidence literature database search for mitigation studies

The Web of Science searches were complemented and expanded by using the Conservation Evidence (https://www.conservationevidence.com/) literature database (Conservation Evidence, 2021). The Conservation Evidence database includes publications of conservation interventions, compiled using systematic searches of both English and non-English language journals (all titles and abstracts) and report series (‘grey literature’) (Sainsbury et al. 2021). To date, systematic searches of over 330 English language journals, over 300 non-English language journals (from 16 different languages) and 24 report series have been conducted (Supplementary material S2 data available at doi: 10.17605/OSF.IO/KQ573). At initial screening, all articles that measured the effect of an intervention that might be done to conserve biodiversity, or that might be done to change human behavior for the benefit of biodiversity were included. English language articles relevant to any reptile species were then read in full and reassessed based on whether the effectiveness of an action to mitigate the impact of dams on freshwater turtles was included. For non-English language articles that passed the initial screening, keyword searches for the terms ‘turtle’ or ‘terrapin’ were carried out, and the title and abstract of the resulting articles were read to check for any mention of freshwater turtles and dams.

### Study data extraction

The following information was extracted from the 47 selected articles: study country, duration (in years), dam function, turtle species and life-stage. Species’ taxonomy, distribution (temperate or tropical latitude) and threat status were obtained from published literature (Rhodin et al., 2018; TTWG et al., 2017). Life-stage was grouped into three classes based on life history and management relevance: early (nest/egg/hatchling), juvenile and/or adult turtle (Lovich et al., 2018; Rachmansah et al., 2020; Shine & Iverson, 1995). Dam function was used to provide an understanding of the representativeness of the selected articles and was not included in the analysis. Function of the dams was obtained from the articles and classified as water supply (including for example irrigation, agriculture, industrial cooling and recreation), hydropower (electricity generation), navigation (transport) and mixed when dams provided multiple functions.

All articles were classified into thematic areas (Table 1) based on the anthropic threats identified in the literature (Alho, 2011; Athayde et al., 2019; Lees et al., 2016; Winemiller et al., 2016). “Solutions” follow the six priority actions for the recovery of freshwater biodiversity identified by Tickner et al. (2020)). For each article we identified a) Threats caused by changes resulting from dams that generated direct or indirect impacts on freshwater turtles; b) Impacts, refers to consequences of these threats; c) Solutions, mitigation actions used or proposed to minimize dam development impacts on freshwater turtles (Table 1).

**Table 1.**
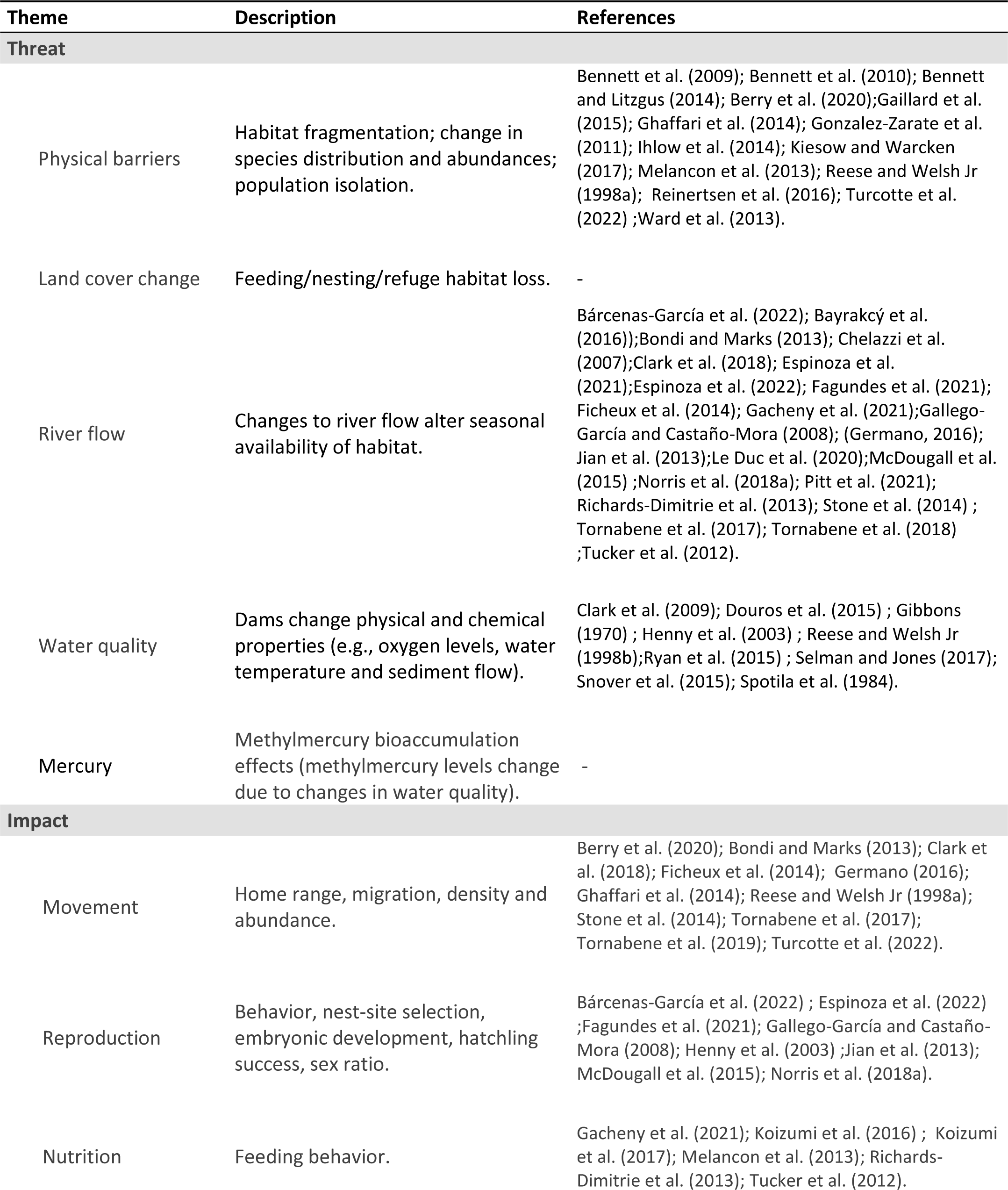
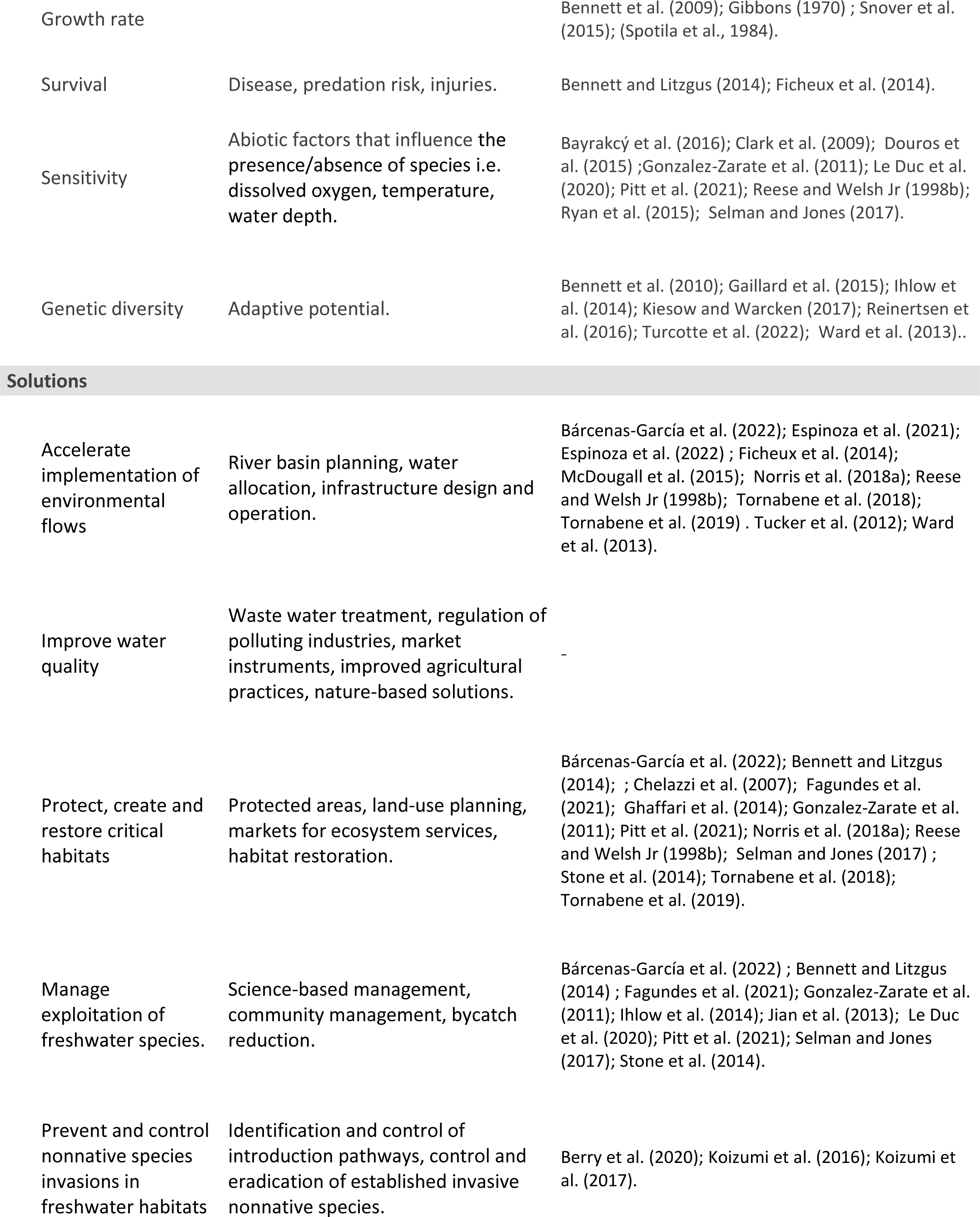
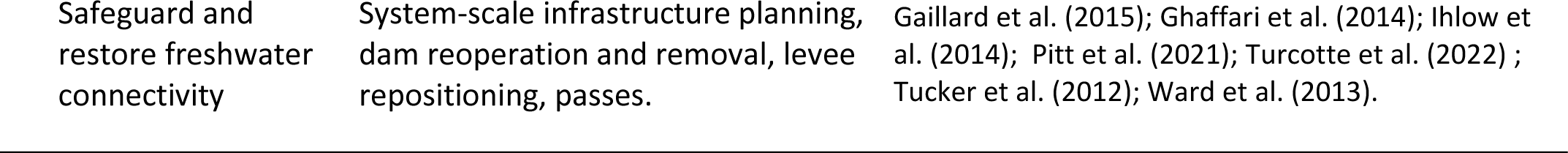
Thematic areas. Thematic areas and typologies used to classify the selected studies. Theme and associated descriptions based on previous reviews of dam impacts (Wu et al., 2019; Zarfl et al., 2019). “Solutions” follow priority actions for the recovery of freshwater biodiversity (Tickner et al., 2020). “References” presents the list of selected studies from the literature search of dam impacts on freshwater turtles, where “-” indicates themes with no studies.

### Data analysis

Patterns in the geographic distribution of publications were evaluated using maps and descriptive statistics. Taxonomic representativeness was assessed using non-parametric tests to compare frequency distributions of studied species with that of extant species per Family (TTWG et al., 2017; Uetz et al., 2021). To understand if studied species could be considered as reflecting 21^st^ century threats, the threat status of studied species was compared against the distribution of extant species (Rhodin et al., 2018). Non-parametric tests were preferred as they are robust and widely adopted for cases with discrete data and small group sizes (Agresti, 2012) and to avoid increased probability of type I errors with parametric frequentist or Bayesian options (Kelter, 2021). Finally, we qualitatively synthesized the effect level on each turtle life-stage as positive (with an ecological or biological benefit); negative (harms the turtle life-stage); and unstudied (if we did not find literature to support it). All analyses were performed in R (R Development Core Team, 2020) with functions available in base R and “tidyverse” collection of packages (Wickham et al., 2019).

## Results

### Geographic and taxonomic bias in the literature

The 47 selected articles included studies based on field surveys (76.6%, n = 36), laboratory research (14.9%, n = 7), interviews (4.3%, n = 2) and modelling inference (4.3%, n = 2). The first article was published in 1970 (Table 1) and measured variation in the reproduction of the pond slider (*Trachemys scripta*) in a reservoir receiving heated effluent from a nuclear reactor in South Carolina, USA (Gibbons, 1970). Only five studies that fitted the selection criteria were published before 2006 and the majority of studies were published during the last ten years, with 74% (n = 32) published between 2012 and 2021 (Table 1). Several studies (17.0%, n = 8) were conducted along waterways with dams providing multiple functions e.g. a mix of water supply, flood control, hydropower, recreation and navigation. Most studies evaluated more localized impacts of dams with single main functions, with water supply/irrigation dams evaluated in 42.6% (n = 20) of selected articles; whereas 38.3% (n = 18) involved hydropower dams and one study was from a predominantly navigational waterway with locks and dams (Berry et al., 2020).

There were clear geographic differences in the number of studies (Fig. 2), with more than half of studies from North America (n = 26), followed by Australia (n = 7). Most studies (72.3%, n = 34) were conducted in temperate latitudes and no studies were found from Africa (Fig. 2). Studies of all three life-stages (early, juvenile and adult) were found from only four countries (Australia, Colombia, Greece and USA, Fig. 2) and only two studies included all three life-stages of the same species (Chelazzi et al., 2007; Gallego-García & Castaño-Mora, 2008). Adult turtles were the main study focus, with 33 articles from 11 countries examining adults (Fig. 2). Additionally, 21 articles from nine countries examined juveniles and nine articles from five countries studied early stages (nests, eggs or hatchling, Fig. 2). Four studies focusing on genetics did not specify the life-stage from which tissue samples were collected (Gaillard et al., 2015; Ihlow et al., 2014; Kiesow & Warcken, 2017; Turcotte et al., 2022). One study evaluated turtle presence and absence at different sites without specifying life-stage (Gonzalez-Zarate et al., 2011) and two studies did not specify the life-stage of captured turtles (Clark et al., 2018; Stone et al., 2014).

**Figure 2.**
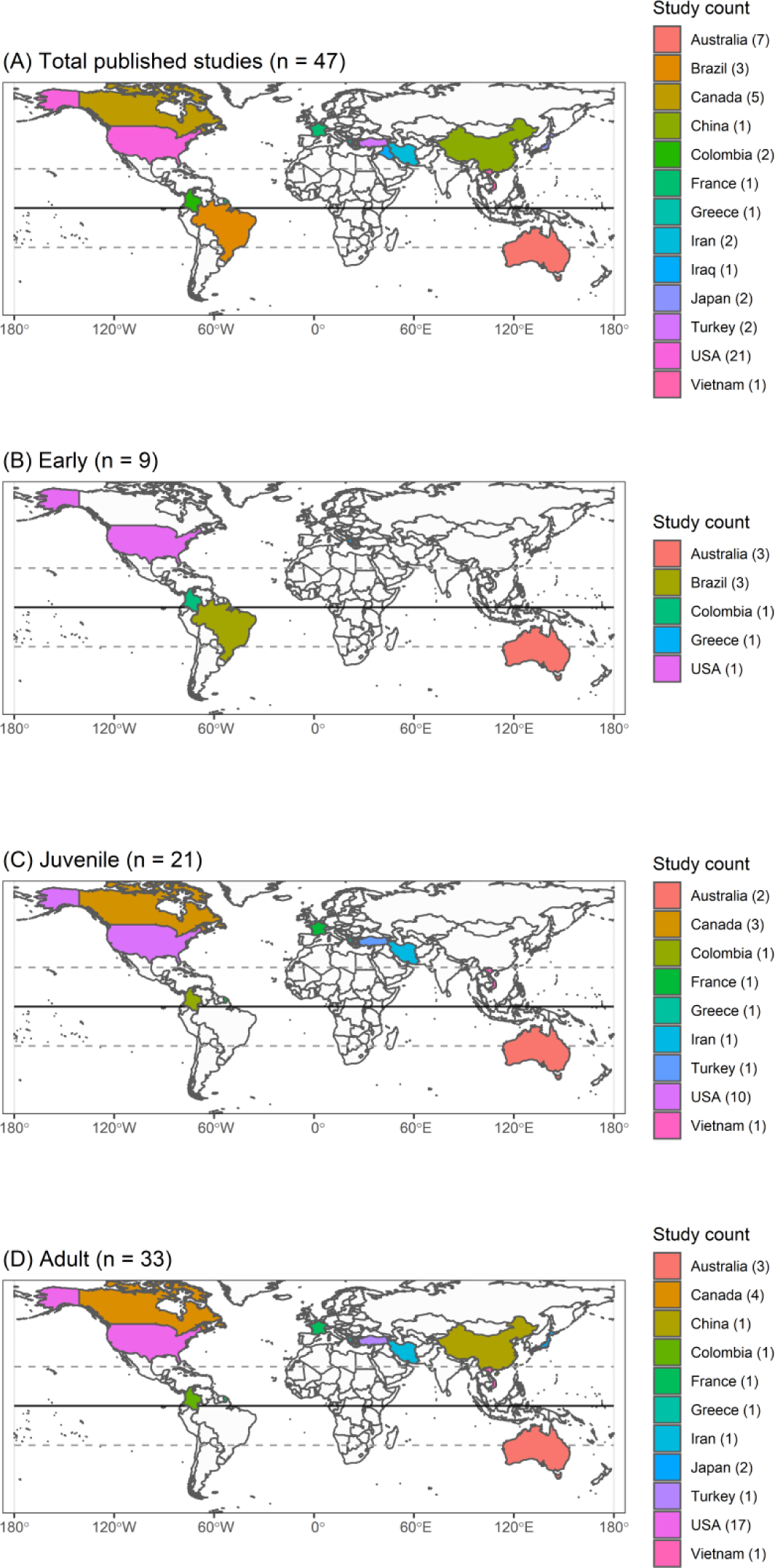
Geographic distribution of articles. Maps showing the geographic distribution of countries where studies were conducted. Showing (A) overall distribution of studies, and countries with studies examining (B) early (nest/egg/hatchling), (C) juvenile or (D) adult life-stages for measures of dam impacts.

Studies examined dam impacts on 30 freshwater turtle species from seven of 11 families (Table 2). Although there was a weak positive relationship, the number of studies was not significantly correlated to the number of extant species in each family (Spearman’s Rho = 0.44, p = 0. 328, Fig. 3). More than a third of studies (42.6%, n = 20) focused on nine North American species of the Emydidae. The Chelydridae (2.1%, n = 1) was the least studied family and Geoemydidae most underrepresented (Fig. 3) relative to extant aquatic turtle diversity (Rhodin et al., 2018; Uetz et al., 2021). The number of species studied in each family was also not significantly correlated to the number of extant species in each family (Spearman’s Rho = 0.36, p = 0.324) and followed a similar pattern to number of studies, with Emydidae species most frequently studied and Chelydridae the most understudied (Fig.3). Of the four unstudied families three included few (5 or fewer) species, but with 27 extant species Pelomedusidae (expected range of 4 to 7 studies, Fig. 3) was the most underrepresented of all families.

**Figure 3.**
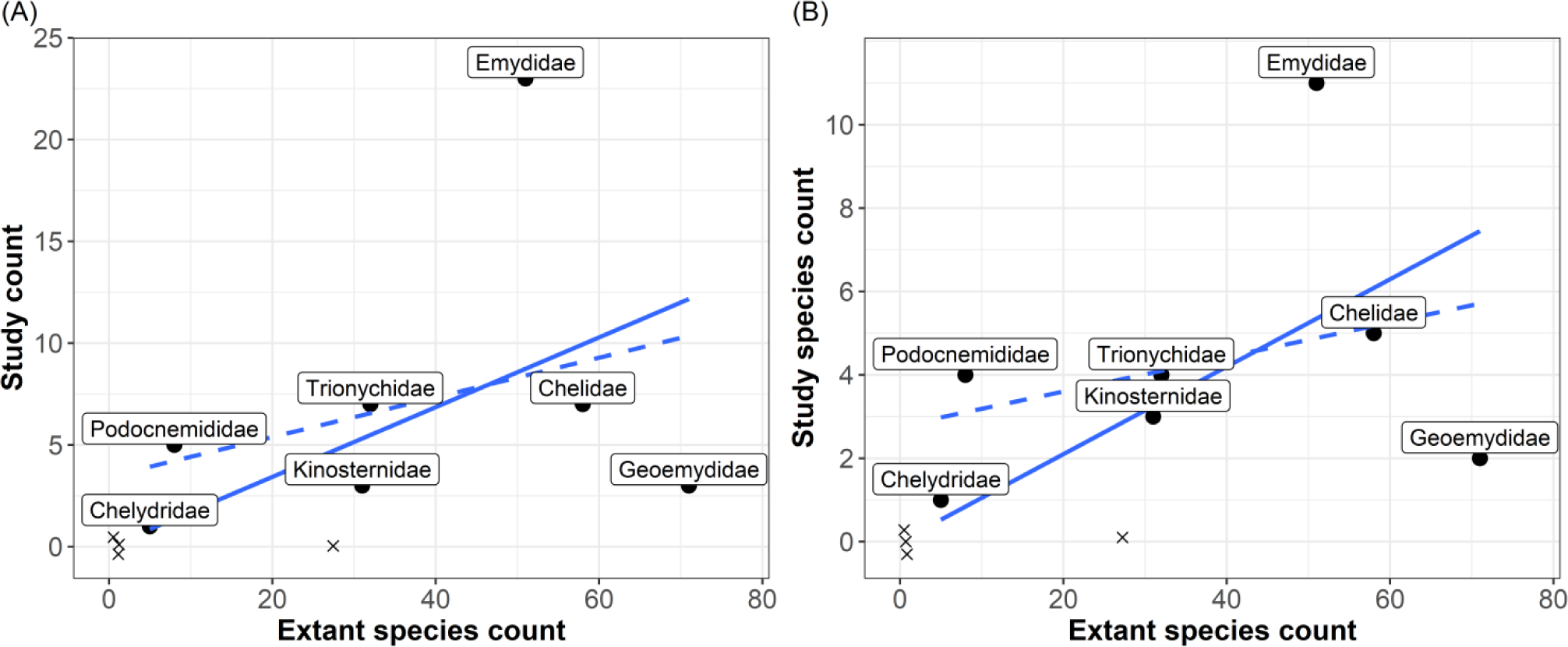
Taxonomic representativeness of articles. Comparison of the number of studies (A) and studied species (B) and extant turtle species per Family [from TTWG et al. (2017))]. Solid lines from a linear model of the expected number in proportion to extant species count, dashed lines from linear model of values obtained from the literature review (lines added to aid visual interpretation). Exes (“x”) show the number of extant species in families with no studies (not included in the linear models, exes are dodged to avoid overlapping).

**Table 2.**
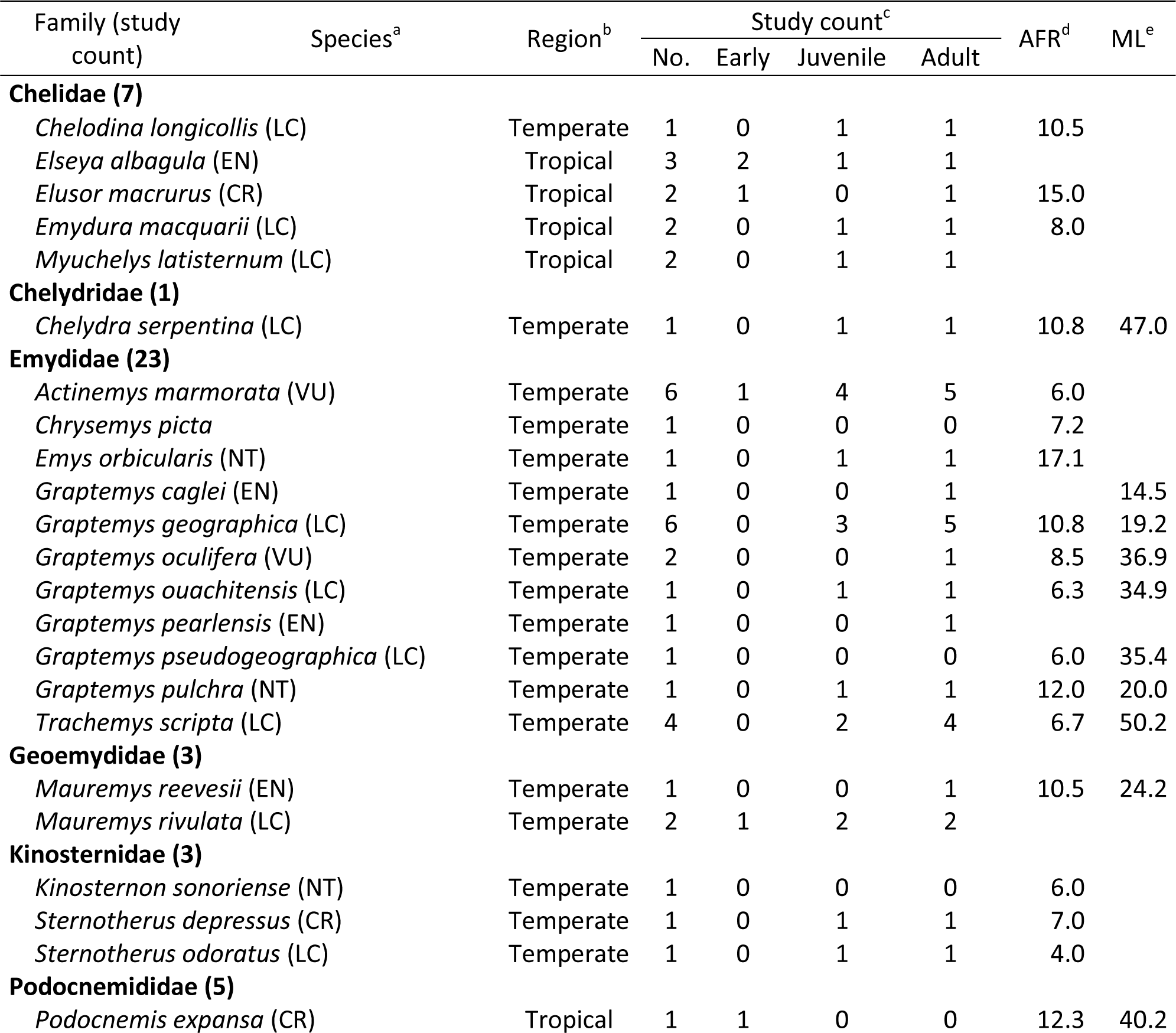
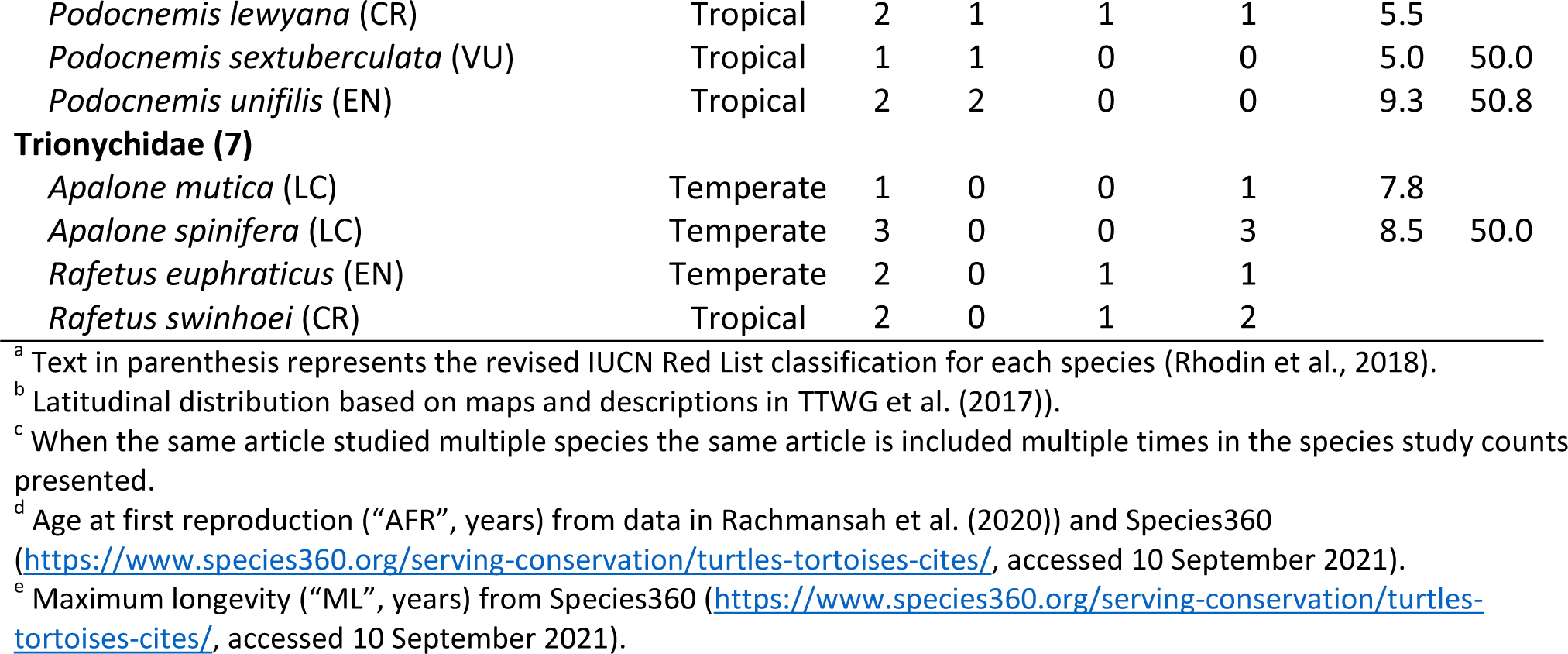
Turtle species. Number of studies examining threats of dams to freshwater turtles obtained from the published literature. Mean values for age at first reproduction (“AFR”, in years) and maximum longevity (“ML”, in years).

Nearly half of the studied species 47% (n = 14) were classified as threatened (CR, EN or VU, Table 2). Whereas 43% (n = 13) of all studied species were classified as Least Concern (LC) and 10% (n = 3) as Near Threatened. The distribution of threatened and nonthreatened species was not significantly different from 50:50 (χ^2^= 0.13, df = 1, p = 0.715) and follows the expected distribution (χ^2^ = 1.44, df = 1, p = 0.230) of the threat status from 360 Testudine species [n = 187 and 138 threatened and unthreatened species respectively, (Rhodin et al., 2018)]. The number of temperate and tropical species studied did differ between threat status categories (Fisher’s Exact Test, p = 0.007), with studies of temperate species having a greater proportion of Least Concern (52 and 22% of studied species, temperate and tropical respectively) and tropical a greater proportion of Critically Endangered species studied (5 and 44% of studied species, temperate and tropical respectively).

Many more studies in temperate regions focused on older age classes (4.2, 37.5 and 58.3% for early, juvenile and adults stages respectively) compared to tropical regions (46.7, 20.0, and 33.3% for early, juvenile and adults, respectively, Fisher’s Exact Test p = 0.0005, Fig. 4). Most studies were of short survey duration, with 70.2% (n = 33) of studies five or fewer years. The vast majority of studies were much shorter than either mean maximum longevity or age at first reproduction of the studied species (Fig. 4). Indeed, there were only seven (14.9%) long-term studies (studies of more than 10 years), all from temperate regions (Fig. 4), with only one study (Pitt et al., 2021) continuing for longer than the maximum longevity of the studied species. There were no long-term studies in tropical regions with the majority of tropical studies focusing on early life-stages (n = 7), whereas studies in temperate regions focused more on juvenile (n = 18) and adult (n = 28) stages (Fig. 4).

**Figure 4.**
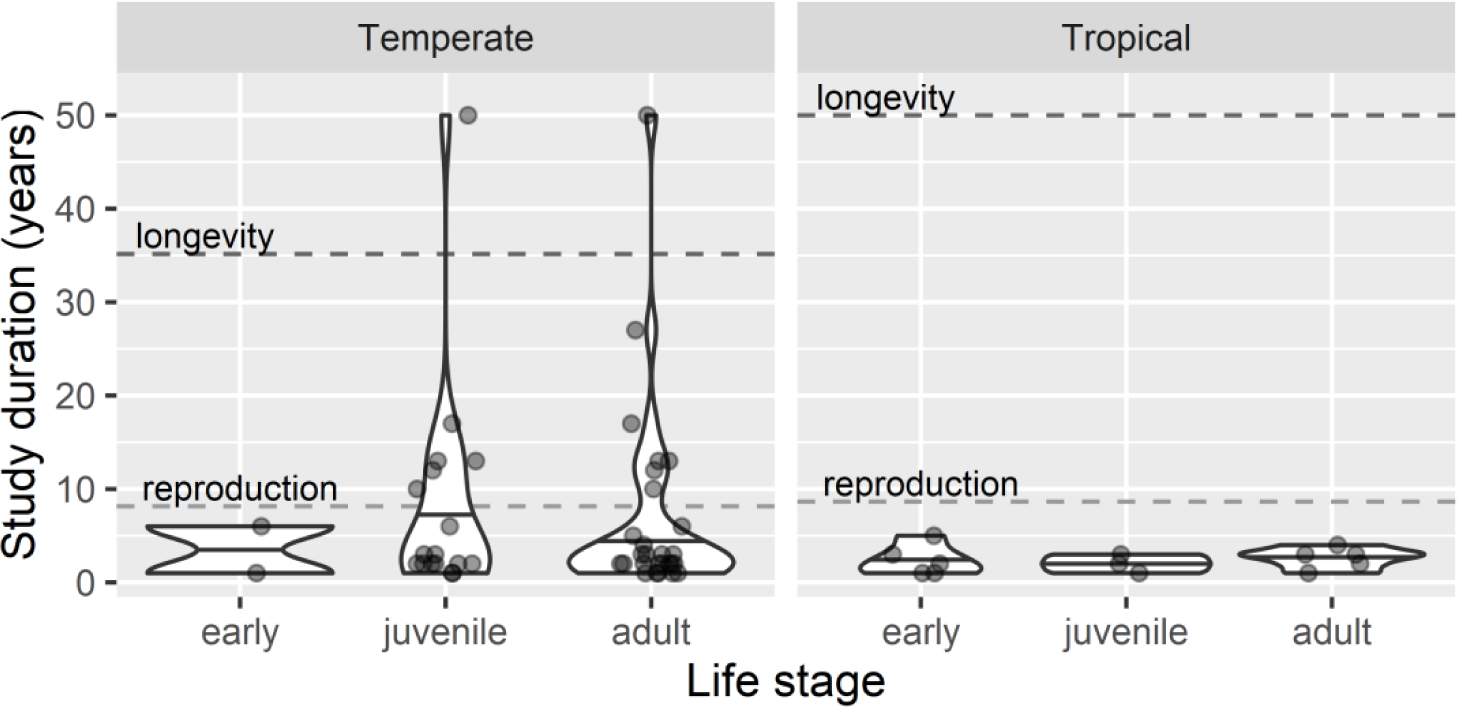
Study duration. Comparison of the years of study examining dam impacts on freshwater turtles in temperate and tropical regions. Distribution of values compared across three life-stage classes (early, juvenile and/or adult, n = 37 studies). When the same article studied multiple life-stages it is included multiple times in the counts presented. Solid horizontal lines within violin plots are 50% quantile of values per life-stage class. Dashed horizontal lines are median values for age at first reproduction (n = 16 and 6 species, temperate and tropical respectively) and maximum longevity (n = 10 and 3 species, temperate and tropical respectively).

### Threats and impacts

A qualitative synthesis of the effects on turtle life-stage (Table 3, Fig. 5) revealed that threats and impacts differed across life-stages and between species. For early stages, changes in river flow and water quality were identified as threats. Indeed changes in river flow were identified as threats across all stages (Table 3). In the juvenile stage, changes in river flow and presence of dams as physical barriers were threats (Melancon et al., 2013). While changes in water quality could also provide potential benefits, this varied between turtle species studied (Clark et al., 2009; Selman & Jones, 2017; Snover et al., 2015). In the adult stage changes to river flows was a threat, whereas changes in water quality and the presence of permanent water provided by physical barriers could also provide benefits for some species (Stone et al., 2014). Changes in land cover and mercury/methylmercury caused by dams remained unstudied in all life-stages.

**Figure 5.**
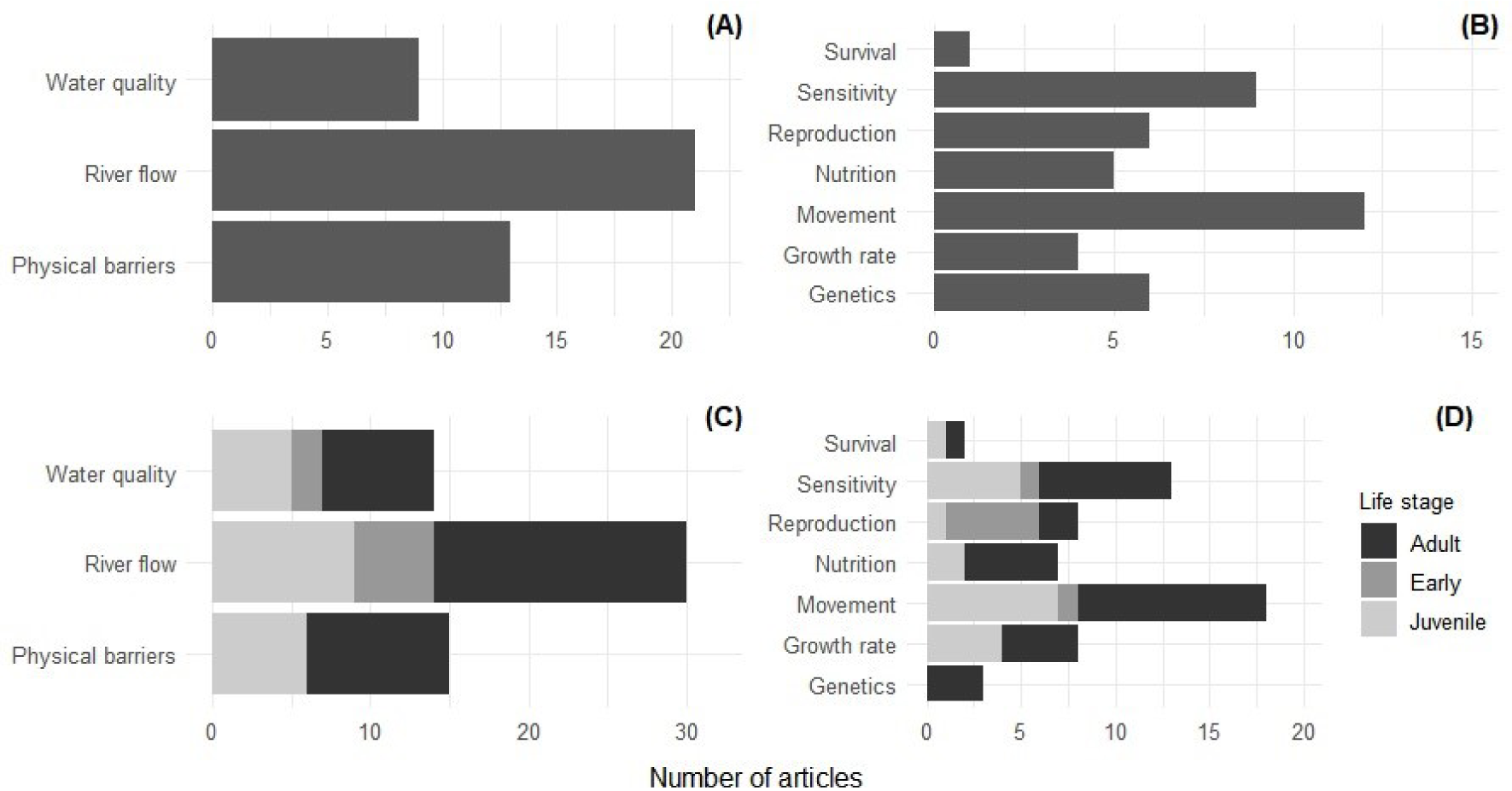
Thematic areas. Total of studies by threats (**A**) and impacts (**B**) and the turtle life-stage (**C** and **D**) according to the thematic areas identified in the review of selected articles. When the same article studied multiple life-stages the same article is included multiple times in the counts presented (C and D).

**Table 3.**
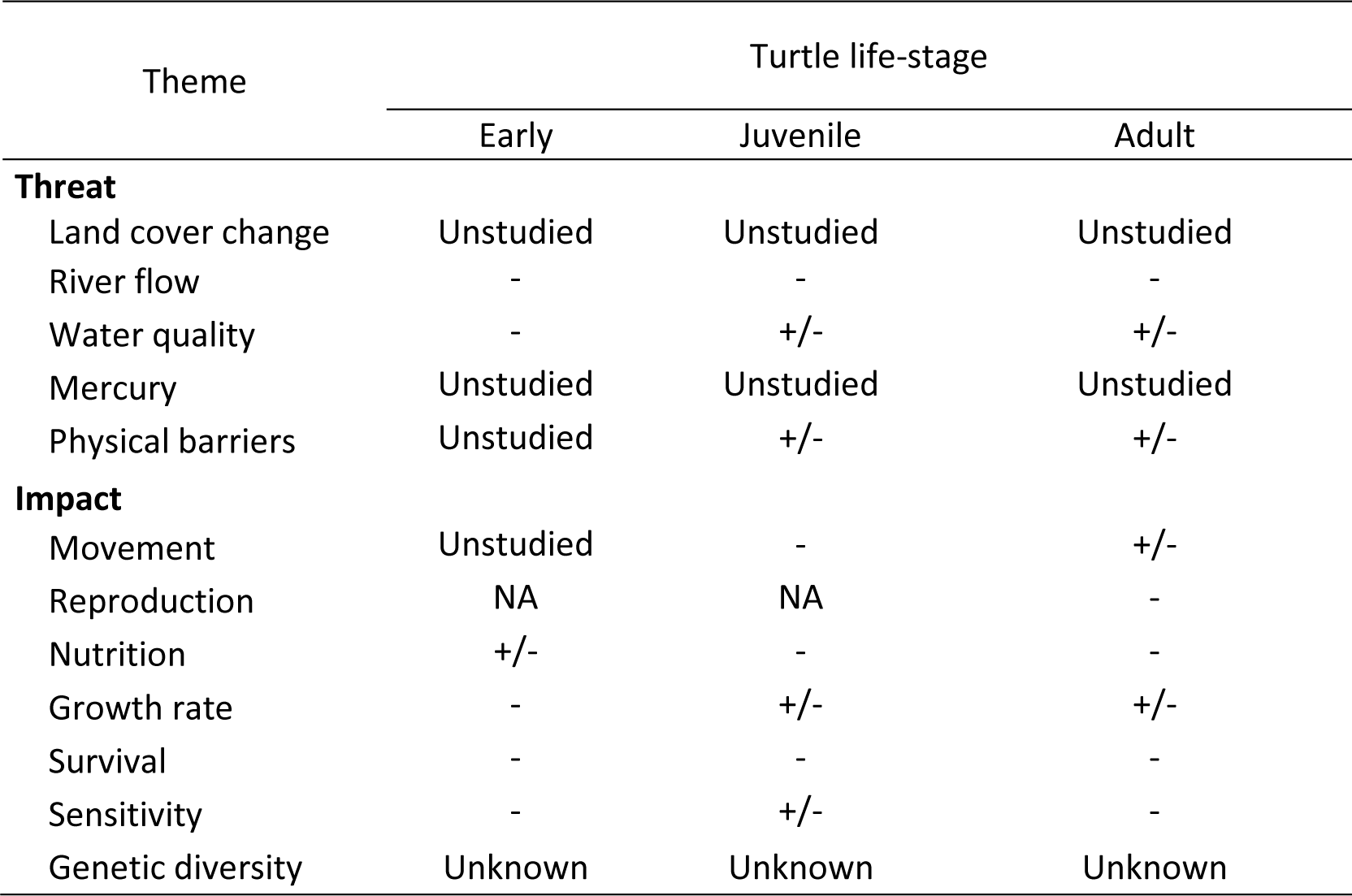
Impact level (negative, positive, unknown or unstudied) from each threat identified and associated dam impacts on freshwater river turtles. “NA” used where the theme is not relevant for the stage i.e. reproduction and early/juvenile stages.

| Theme | Turtle life-stage |  |  |
| --- | --- | --- | --- |
|  | Early | Juvenile | Adult |
| **Threat** |  |  |  |
| Land cover change | Unstudied | Unstudied | Unstudied |
| River flow | - | - | - |
| Water quality | - | +/- | +/- |
| Mercury | Unstudied | Unstudied | Unstudied |
| Physical barriers | Unstudied | +/- | +/- |
| **Impact** |  |  |  |
| Movement | Unstudied | - | +/- |
| Reproduction | NA | NA | - |
| Nutrition | +/- | - | - |
| Growth rate | - | +/- | +/- |
| Survival | - | - | - |
| Sensitivity | - | +/- | - |
| Genetic diversity | Unknown | Unknown | Unknown |

The impacts also differed across life-stages and between species, but when studied, negative impacts were documented for all life-stages and all themes (Table 3). In the juvenile stage, there could be positive impacts on growth rate and sensitivity, for example as new environmental conditions may create refuge habitat for juveniles (Ryan et al., 2015). In adults the creation of waterways could facilitate movements, range expansions and exchanges between once isolated species (Berry et al., 2020). The impacts on genetic diversity were not differentiated across different life-stages, but studies documented evidence of negative impacts of dams on genetic diversity (Ihlow et al., 2014; Turcotte et al., 2022).

### Mitigation actions

A total of five studies tested the effect of four mitigation actions [Fig. 6, (Espinoza et al., 2021; Espinoza et al., 2022; Ficheux et al., 2014; Pitt et al., 2021; Stone et al., 2014)]. Of these studies, three were from temperate regions with long-term data collection spanning 50 (Pitt et al., 2021), 20 (Stone et al., 2014) and 17 years (Ficheux et al., 2014) and two were relatively short term 3 year studies from sub-tropical Australia (Espinoza et al., 2021; Espinoza et al., 2022). One article (Pitt et al., 2021) documented the effect of dam removal on the northern map turtle (*Graptemys geographica*). A study from the USA (Stone et al., 2014) demonstrated the potential of volunteers to help implement actions (dam repair and silt removal) to maintain artificial impoundments for the Sonora mud turtle (*Kinosternon sonoriense*). Another from southern France (Ficheux et al., 2014) showed how earlier flooding across wetland areas improved hibernation success for the European pond turtle (*Emys orbicularis*); whilst studies from Australia (Espinoza et al., 2021; Espinoza et al., 2022) demonstrated how environmental flow management could facilitate movements of adult Mary River turtles (*Elusor macrurus*) and improve nesting success for the Endangered white-throated snapping turtle (*Elseya albagula*).

**Figure 6.**
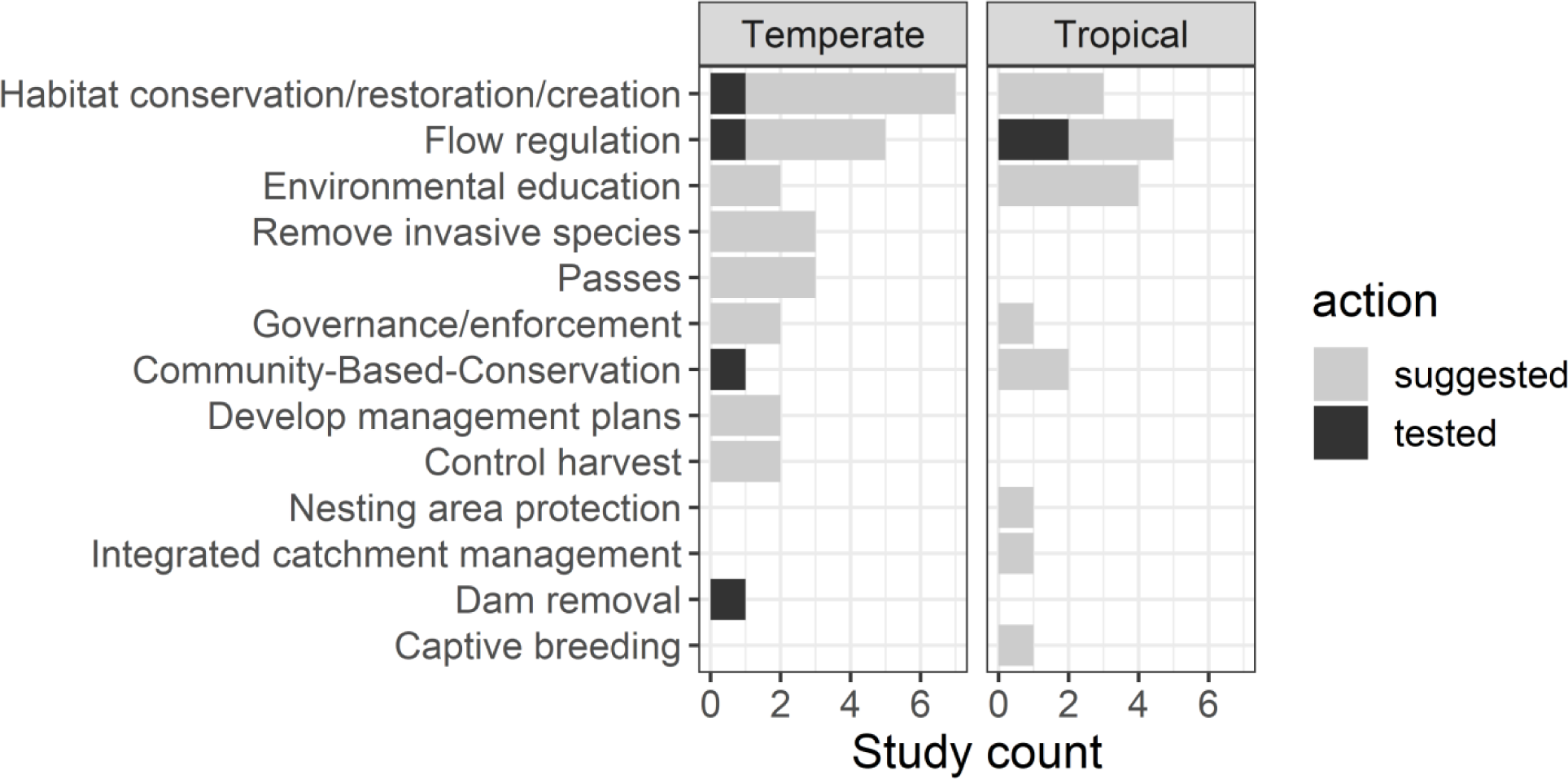
Mitigation actions. Actions presented in 27 of 47 selected articles compared between studies of freshwater turtle species in temperate and tropical regions.

Far more studies (57%, n = 27) presented suggestions rather than tested possible conservation actions. The principal suggestions to mitigate dam impacts (Fig. 6) were habitat conservation/restoration/creation (9 studies), environmental education (6 studies) and flow regulation (6 studies). While improving governance and enforcement was suggested three times [improved regulation of recreational boating (Bennett & Litzgus, 2014), more rigorous environmental impact assessments (Norris et al., 2018a) and additional state/federal protections for declining species (Selman & Jones, 2017)].

## Discussion

To our knowledge, our study constitutes the most extensive search of the global scientific literature for assessing the impact of dams on freshwater turtles and mitigation measures that have been carried out to date. The inclusion of over 300 non-English language journals increases confidence that important sources of evidence for actions to mitigate impacts have not been missed (Amano et al., 2021). This comprehensive systematic review of existing evidence provides insight not only to the trends, thematic fields and gaps in current research but also conservation solutions to the impacts caused by dams on freshwater turtles. Studies were primarily focused on river flow changes, however, there were few studies in some regions, principally the tropics, and gaps on important themes like bioaccumulation of methylmercury linked to dams. We first discuss geographic and taxonomic biases, then the impacts of dams on freshwater turtles. Finally, we describe the mitigation actions proposed and implemented for turtle recovery and conservation.

### Geographic and taxonomic bias

There were marked differences in the scientific production between temperate and tropical latitudes. Our finding that 62.5% of studies were from USA and 66% of the total studies were from temperate latitudes confirms results from a previous study that showed most scientific knowledge came from temperate regions (Rachmansah et al., 2020). This geographic bias was particularly surprising as tropical regions have a high potential for hydropower development (Grill et al., 2019) and are also considered priority areas for turtle conservation (Mittermeier et al., 2015; Stanford et al., 2020). There was also a connection between geographic biases and study taxa as most studies (42.6%) focused on ten North American species of the Emydidae and the lack of studies from Africa meant the Pelomedusidae (freshwater turtles native to sub-Saharan Africa) was not represented.

It is possible that the lack of studies on impacts from Africa and Asia could be a result of the English language Web of Science searches. Based on evidence from recent studies it does however appear likely that there are indeed very few studies documenting impacts on freshwater turtles across African and Asian basins impacted by dams. For example a recent special issue regarding the Lower Mekong basin included no articles examining freshwater turtles (with 20 published articles, https://www.mdpi.com/journal/water/special_issues/Mekong_River#published, accessed 8 September 2021). The inclusion of the Conservation Evidence database with its comprehensive coverage of both English and non-English language literature increases confidence that the geographic patterns are most likely a result of lack of documented evidence and not language based search bias (Amano et al., 2021).

### Threats and impacts of dams on freshwater turtles

Based on our results, the principal changes that impacted freshwater turtles were river flow modification, physical barriers and water quality. Land cover change was not assessed as a direct impact of dams on freshwater turtles.

### Changes in river flow

#### Loss of feeding habitat

Changes in the flood pulse together with the loss of feeding habitat can strongly impact the availability of food resources (Bennett et al., 2009; Petrov et al., 2018). This was shown in Australia where the diet of three freshwater turtle species was compared in sites with and without damming impact (Tucker et al., 2012). Damming reduced ingestion of subaquatic plants and fruits by *Emydura kreffti* and *Elseya albagula* and *Myuchelys latisternum* showed a diminished ingestion of aquatic invertebrates in impacted habitats (Tucker et al., 2012). The changes in diet may reflect changes in availability of resources associated with damming, as a study showed increased consumption of bryozoans (including Zebra mussels) in a reservoir compared with previous studies from downstream before Zebra Mussel establishment. The flood pulse also plays a role in access to food. For example, insect larvae may depend on shallow waters for their development, and macrophytes are also reduced when water levels fluctuate (Tucker et al., 2012). Therefore, the adaptive response of turtles to dams may depend on their capacity to change foraging strategies (Petrov et al., 2018; Richards-Dimitrie et al., 2013).

#### Loss of nesting habitat

Reproduction of freshwater turtles can be tied to the seasonal availability of nesting habitats. Nests of turtle species that use terrestrial nesting habitats (e.g. river banks) may be at greater risk from flooding (Bodie, 2001), as is the case of species of the South American genus *Podocnemis* (Eisemberg et al., 2016; Gallego-García & Castaño-Mora, 2008; Norris et al., 2018a; Norris et al., 2020), genus *Elseya* in Austrailia (Espinoza et al., 2022), and of the North American Emydidae and Trionychidae (Pitt et al., 2021; Tornabene et al., 2018). Changes to a river’s annual discharge cycle caused by dams can therefore reduce the availability of nesting areas (Bárcenas-García et al., 2022; Norris et al., 2018a; Tornabene et al., 2018), as well as the duration of low water levels, which may affect the behavior and nesting success of these species (Eisemberg et al., 2016; Espinoza et al., 2022; Tornabene et al., 2018).

The artificial regulation of dammed rivers can result in permanent flooding and/or a reduction in nesting areas. In the absence of adequate nesting habitat, nest-site selection by females may be compromised as nests may be placed even if they may not represent a good choice for females, eggs or hatchlings (Boyer, 1965; Kolbe & Janzen, 2002; Refsnider & Janzen, 2010; Schlaepfer et al., 2002). Nest-site selection can affect the nest’s vulnerability to predation by wildlife (Spencer, 2002) and humans (Michalski et al., 2020) and may even render nests susceptible to submersion by flash floods that can occur due to hard-to-predict events, resulting from climate change (Eisemberg et al., 2016), or inflow impoundment by dams (Espinoza et al., 2022; McDougall et al., 2015; Norris et al., 2020). Water released by the Kota hydropower dam in India caused the Chambal River to rise, flooding nesting areas and causing loses of 7.7% and 9.6% of the nests of *Batagur kachuga* and *Batagur dhongoka* respectively (Rao & Singh, 1987). In Brazil, the filling of a hydropower dam reservoir resulted in the flooding and loss of 3.9 hectares of nesting habitats and areas used by the yellow-spotted river turtle *Podocnemis unifilis* (Norris et al., 2018a). Besides Norris et al. (2018a) and (Bárcenas-García et al., 2022) who applied before-after control-impact study design, no other study evaluated freshwater turtle nesting patterns with baseline monitoring previous to dam installation.

Changes in nest microclimate, for example changes in substrate humidity and/or temperature, can also affect sex ratio, embryonic development and hatching success (Refsnider et al., 2013). Eggs of species adapted to nesting on land may withstand brief flooding [e.g. up to two days for *Podocnemis unifilis* eggs (Norris et al., 2020)] but permanent immersion during the early stages of incubation diminishes embryo survival (Bodie, 2001). Indeed, eggs of Chelidae *Emydura krefftii* may not tolerate being under water for more than half an hour (Hollier, 2012). Barriers created by dams also limit transportation and downstream availability of nutrients and sediments. This change drastically reduces the volume of sediment that can be transported downstream leading to the progressive disappearance of potential nesting areas for freshwater river turtles (Le Duc et al., 2020; Lenhart et al., 2013).

### Physical barriers

The fragmentation of free flowing river habitat limits migrations, causes isolation and diminishes the genetic flow between populations of freshwater turtles (Gallego-García et al., 2018). Damming divides populations, making reproduction more difficult (Jian et al., 2013) and decreasing the adaptive capacity of impacted populations as genetic diversity is lost. This could result in inbreeding, with potential consequences for reproductive fitness and survival in the disturbed environment (Turcotte et al., 2022). These are factors that, together, have implications on population recruitment (Bennett et al., 2010; Buchanan et al., 2019; Gallego-García et al., 2018; Ihlow et al., 2014). Evaluating impacts of dams as barriers at a genetic level can however take several generations and depending on the species can require many decades if not centuries for the changes to manifest (Bennett et al., 2010; Gaillard et al., 2015; Kiesow & Warcken, 2017; Reinertsen et al., 2016; Turcotte et al., 2022; Ward et al., 2013).

In Canada, the dispersal and occurrence of *Graptemys geographica* females declined in areas fragmented by locks, dikes and hydropower dams (1.53 ± 0.31 km), compared to females found in contiguous areas [8.51 ± 1.59 km, (Bennett et al., 2010)]. The abundance of both generalist (*E. macquarii*) and more specialist (*M. latisternum*) species declined after five years in locations near a dam in Australia (Clark et al., 2018). A decrease in the abundance of fish and absence of the Yangtze giant softshell turtle *R. swinhoei* was recorded after the installation of the Hoa Binh hydropower dam in Vietnam, according to interviews with fishermen (Le Duc et al., 2020). Similarly in China, after sand banks were flooded by the Nansha hydropower dam in 2006, it was no longer possible to detect the presence of *R. swinhoei* (Jian et al., 2013).

Permanently inundated lotic environment created by dams can favor certain species including non-native turtle species (Berry et al., 2020). Damming could provide new potential habitat for freshwater turtles, like permanent impoundments of otherwise ephemeral streams (Stone et al., 2014). Water supply dams for cattle can also be used as permanent habitat by generalist species like *Actinemys marmorata* [Table 3, (Germano, 2016)]. But, such cases depend on active management to maintain healthy populations, as highlighted by an example from Europe showing the importance of managing cattle to avoid trampling and appropriate management of flow regimes for the impacted turtle species (Ficheux et al., 2014).

### Changes in water quality

A lack of oxygen may limit the presence and persistence of diving species in reservoir environments. Lack of oxygen in reservoirs has implications for turtle physiology as it limits the capacity to obtain oxygen from the water, which can reduce diving ability by 51% (Clark et al., 2018). The impact of changing oxygen levels was recorded in Australia for *Elusor macrurus* hatchlings, a species with bimodal respiration which, nevertheless, cannot withstand hypoxia conditions for long periods of time (Clark et al., 2009). Eutrophication may however benefit some turtle species, for example an increase in emergent vegetation could be potential refuge habitat for the juvenile stages of *Chelodina longicollis* [Table 3, (Ryan et al., 2015)]. Additionally, adults of *Trachemys scripta* were benefited by warmer water temperatures in a nuclear reactor cooling reservoir that increased the time available for foraging and increased growth rates (Gibbons, 1970). These increases could be facilitated by behavioral plasticity as the same species also showed behavioral thermoregulatory adaptations e.g. differences in aquatic and atmospheric basking depending on proximity to warmer water (Spotila et al., 1984). In contrast, dammed rivers with cooler temperatures delayed reproductive maturity by 9 years in western pond turtles [*Actinemys marmorata*, (Snover et al., 2015)].

Accumulation of contaminants in dam reservoirs was reflected in the pesticide concentration found in the common snapping turtle (*Chelydra serpentina*), which increased closer to a water supply dam (Douros et al., 2015). Additionally, the new physical-chemical environment (e.g. lower pH) and the elevated rates of decomposition of submerged organic matter can increase mercury methylation by bacteria around dams and reservoirs (Millera Ferriz et al., 2021; Regnell & Watras, 2019). Methylmercury is an extremely toxic contaminant that can be highly damaging to people (Budnik & Casteleyn, 2019) and aquatic vertebrates including turtles (Green et al., 2010; Meyer et al., 2014). As freshwater turtles are often consumed by riverside populations there is a strong potential for freshwater turtles to represent a source of dietary methylmercury (Green et al., 2010). Patterns of mercury and methylmercury contamination and bioaccumulation have been intensely studied around dams (Millera Ferriz et al., 2021; Wang et al., 2004) and in turtles from temperate (Burger & Gibbons, 1998; Meyer et al., 2014; Slimani et al., 2018) and tropical regions (Eggins et al., 2015; Schneider et al., 2009; Schneider et al., 2010). The lack of studies assessing bioaccumulation of methylmercury in freshwater turtles in and around dams was therefore surprising.

### Mitigation actions

Among the measures to mitigate/minimize dam impacts on freshwater turtles the most frequently suggested were habitat conservation/restoration/creation (Ghaffari et al., 2014; Gonzalez-Zarate et al., 2011; Norris et al., 2018a; Pitt et al., 2021; Reese & Welsh Jr, 1998b; Tornabene et al., 2019), flow regulation and environmental education (Ghaffari et al., 2014; Gonzalez-Zarate et al., 2011; Ihlow et al., 2014; Jian et al., 2013; Le Duc et al., 2020; Norris et al., 2018a). Promoting habitat creation and restoration would likely contribute to long-term conservation of breeding and nesting areas, as well as potential foraging areas for turtles. Restored vegetation can also help reduce erosion, improve water quality, and promote the reestablishment of a wide variety of aquatic and terrestrial species (Santoro et al., 2020). However, no published studies were found that evaluated the implementation of these suggested measures for freshwater turtle species impacted by dams.

Studies also suggested that companies in charge of dam development and operation should adopt a holistic vision of catchment management, including measures such as flow regulation for the specific freshwater turtles impacted and adapting inlets to favor turtle dispersal between rivers and the available flood plains (Howard et al., 2017; McDougall et al., 2015). Indeed an example from Australia showed that such flow rate changes could benefit multiple species without negatively affecting water supply to end users (Espinoza et al., 2022; McDougall et al., 2015).

Although more than half of studies suggested mitigation actions, only five evaluated interventions. This pattern follows worrying trends where for threatened species, only a small proportion of available budgets are implemented, with an example from the US demonstrating that only a small fraction of proposed management tasks for species recovery are achieved (Gibbs & Currie, 2012). Although the majority of studies suggested the need for additional research including long-term monitoring, monitoring is not sufficient to solve conservation problems (Buxton et al., 2020; Legg & Nagy, 2006). It is worth noting that a number of other studies were highlighted within the Conservation Evidence database that evaluated a range of interventions with potential relevance to the threat of dams on freshwater turtles. For example, studies evaluating habitat restoration/creation (e.g. “Create or restore ponds”) or education and awareness raising (e.g. “Engage local communities in conservation activities”) may provide evidence that could be applicable for a wide range of taxa and be implemented in response to a large number of threats, including those arising from dams (Conservation Evidence, 2021; Sainsbury et al., 2021). While interventions and actions developed and implemented within local contexts may well have the most relevant results, it remains an open question the extent to which relevant evidence can be shared across different species groups, habitats and contexts. Such sharing of evidence could go some way to filling gaps in the literature and increase the collective capacity for using evidence to inform conservation decision making.

Facilitating movements around dams is likely to help maintain connectivity and reduce mortality. Yet, there is little evidence available for interventions such as passes except for fishes. There are examples of freshwater turtles using fish passes but not necessarily for movements, but rather as feeding locations (Agostinho et al., 2012). Evidence is needed to inform the development of passes that can be effective for both small and large turtles (>30 kg) in rivers with high predator diversity. Another option is implementing habitat modifications to ensure safe terrestrial passage around dams. Although we did not find any evidence for the efficacy of such actions around dams, there are examples of habitat modifications e.g. barrier installation (Heaven et al., 2019), which have also been used together with the creation of suitable nesting habitats to reduce adult mortality around roads (Nagle & Congdon, 2016).

There is an increasing need to develop integrated and adaptive approaches to mitigate dam impacts on freshwater turtles. Community-based Conservation encourages social organization and the creation of initiatives to conserve natural capital. Several studies have already shown that the survival of turtle hatchlings and adults increases through conservation by community management (Campos-Silva et al., 2018; Norris et al., 2020; Norris et al., 2019; Rivera et al., 2021; Stone et al., 2014). Community-based Conservation also encourages participation to monitor, protect, and reduce predation of freshwater turtles within communities (Campos-Silva et al., 2018; Rivera et al., 2021; Vallejo-Betancur et al., 2018). Actions may also include rescue activities such as that which occurred in the eastern Brazilian Amazon, where community-based actions contributed to the rescue of 926 eggs, 65 premature hatchlings and the release of 599 hatchlings of *P. unifilis* during the flooding of nests by rising water levels (Norris et al., 2020). However, community participation in any conservation project requires that the communities are actively involved in creating plans and/or management projects (Campos-Silva et al., 2018; Rivera et al., 2021), procuring sources of economic income, as well as providing the necessary inputs for project development so that they are not abandoned due to lack of resources (Norris et al., 2018b; Stone et al., 2014).

It is necessary to strengthen the protection and monitoring of existing nesting areas (Forero-Medina et al., 2019), and of juvenile and adult stages (Hance, 2020). Such actions should be supported by additional research to establish if population recruitment is occurring and provide more robust estimates of turtle population dynamics particularly in the tropics (Norris et al., 2019; Rachmansah et al., 2020). This could be achieved with the promotion of social, governmental, business and research center participation (Guo et al., 2021). It may also be possible to complement this with environmental education actions at different levels of society (including children), to revalue the importance of freshwater turtles as components of the ecosystem and diverse cultures (Ghaffari et al., 2014; Gonzalez-Zarate et al., 2011; Le Duc et al., 2020). Another strategy would be to implement community management to regain the cultural, economic, ecological, political and social values of the communities over their natural resources (Brownson & Fowler, 2020; Campos-Silva et al., 2018; Harper et al., 2021; Lopes et al., 2021).

In addition, to mitigate dam impacts and prevent the loss of species and ecological functions, environmental authorities must conduct more robust and rigorous Environmental Impact Assessments (Norris et al., 2018a), as well as provide support to supervise compliance with mitigation actions and monitoring effectiveness (Guo et al., 2021; Valiente-Banuet et al., 2015).

### Implications for Conservation

Actions that mitigate known negative impacts are urgently required to prevent the collapse of populations of critically endangered freshwater turtle species. Our review showed that impacts of dams on freshwater turtles remain poorly studied, particularly in tropical regions. Changes in the river flow caused by dams on freshwater turtles were the principal focus, but there were important information gaps regarding the effects of changes in land cover, methylmercury bioaccumulation and water quality. With only five studies evaluating interventions, much more evidence is required to evaluate mitigation actions across different life-stages and geographic regions. Integrated monitoring programs that provide evidence at relevant spatial and temporal scales for all turtle life-stages are needed to promote the conservation of these threatened species.

## Acknowledgements

We would like to thank the Universidade Federal do Amapá for providing logistical support; Andrea Bárcenas received a master scholarship provide by Coordenação de Aperfeiçoamento de Pessoal de Nível Superior (CAPES). The work with Conservation Evidence was funded by Arcadia, The David and Claudia Harding Foundation and MAVA.

## Declaration of competing interest

The authors declare that they have no known competing financial interests or personal relationships that could have appeared to influence the work reported in this paper.

## Notes

### Competing Interest Statement

The authors have declared no competing interest.

### Summary of Updates

Various stylistic changes to address reviewer comments. Literature search updated to April 2022. Text updated to reflect addition of new studies.

